# Mechanics reveals the role of peristome geometry in prey capture in carnivorous pitcher plants (*Nepenthes*)

**DOI:** 10.1101/2023.04.20.537328

**Authors:** Derek E. Moulton, Hadrien Oliveri, Alain Goriely, Chris Thorogood

## Abstract

Carnivorous pitcher plants (*Nepenthes*) are a striking example of a natural pitfall trap. The trap’s slippery rim, or peristome, plays a critical role in insect capture via an aquaplaning mechanism that is well documented. Whilst the peristome has received significant research attention, the conspicuous variation in peristome geometry across the genus remains unexplored. We examined the mechanics of prey capture using *Nepenthes* pitcher plants with divergent peristome geometries. Inspired by living materials, we developed a mathematical model that links the peristomes’ three-dimensional geometries to the physics of prey capture under the laws of Newtonian mechanics. Linking form and function enables us to test hypotheses related to the function of features such as shape and ornamentation, orientation in a gravitational field, and the presence of ‘teeth’, while analysis of the energetic costs and gains of a given geometry provides a means of inferring potential evolutionary pathways. In a separate modeling approach, we show how prey size may correlate with peristome dimensions for optimal capture. Our modeling framework provides a physical platform to understand how divergence in peristome morphology may have evolved in the genus *Nepenthes* in response to shifts in prey diversity, availability, and size.

**Significance Statement:** Pitcher plants (*Nepenthes*) produce an astonishing array of leaf-derived traps into which preys (typically insects) slide from a rim (the peristome). How prey capture varies across this varied genus is a mystery. We hypothesized that insects behave differently depending on peristome size and geometry. We examined the physics of prey capture under the laws of Newtonian mechanics to show that prey size and behavior correlate with peristome shape. We conclude that a diversity of peristomes in *Nepenthes* evolved in response to variation in prey capture.

---

Carnivorous plants evolved various forms of leaf-derived traps that attract, capture, retain, kill, and digest animal prey, as a mode of survival in nutrient-poor environments. *Nepenthes* is a tropical genus of carnivorous pitcher plants that produce specialized pitfall traps. Insects are attracted by lures such as coloration and nectar, and become trapped when they ‘aquaplane’ off the slippery pitcher rim (peristome), a surface structured with specialized ridges (1, 2), leading them to fall into a vessel of digestive fluid (3). The insects release nitrogen which gives the plants a strong selective advantage in environments where light and water are plentiful, but nutrients are limiting (4).

The specialized trapping surfaces of carnivorous *Nepenthes* pitcher plants are receiving growing interest from biologists and engineers because of their strong biomimetic potential (5). For example, the slippery trapping surface of the *Nepenthes* pitcher has inspired Slippery Liquid-Infused Porous Surfaces (SLIPS) which have exceptional wettability performance (6, 7). Yet despite research focused on the peristome as a key feature in the evolution of the trap, and as a source of inspiration to technologists, little is known about the mechanics of prey capture in *Nepenthes*, or how this varies among species.

To date, there are 179 accepted species of *Nepenthes* (POWO, 2022) and they show an astonishing diversity in pitcher morphology. Little is known about the prey trapped by most species in nature. Among the few species in the genus examined, diversity seems to mirror a range of nutrient acquisition strategies linked to habitat characteristics (8). For example, ants are a common form of prey in lowland habitats (9), whereas flying insects are often trapped by plants growing in mountain environments (10). More specifically, research in the last two decades has revealed that divergent pitcher morphology is linked to nutrient acquisition sources ranging from termites (9), and leaf litter (11), to mammalian feces (12, 13). Most recently, a species was reported from Borneo that produces pitchers underground (14). This diversity in pitcher function appears to be the result of an adaptive radiation driven by dietary shifts, analogous to well-known examples in animals, such as the diverse beak shapes of Darwin’s finches and the various adaptations of cichlid fish in the African Great Lakes (3). However, only a fraction of the diversity of *Nepenthes* has been examined, and we know little or nothing of the prey spectrum for most species.

The general mechanism by which insects slide off the *Nepenthes* is well documented. A film of water stabilizes on the superhydrophilic surface (1). The surface is covered by a regular, hierarchical microstructure of parallel ridges, or channels (2, 5). These ridges guide prey into the trap in a controlled, non-arbitrary way (5). Recent work shows that macroscopic ridges restrict lateral but enhance radial spreading of water, creating slippery chutes. Meanwhile, microscopic ridges ensure the watery film between the insects’ feet and the peristome remains stable, causing insects to aquaplane (2). These principles seem to be consistent across multiple species, indicating a common mechanism underlying insect aquaplaning. However, the gross morphology of peristomes is conspicuously diverse, ranging from cylindrical rims to highly ornate, fluted, and toothed structures (Fig. 1). Disparate geometries in peristomes could be linked to ecological niche. For example *N. veitchii* [Fig. 2(c)] has an unusual life history: the plant clings to trees with the pitchers oriented such that the ventral surface is parallel to the tree surface. Meanwhile species such as *N. macrophylla* and *N. diabolica* [Fig. 2(d)] produce pitchers, often half-buried in moss, with conspicuously toothed peristomes. The prey spectrum of these, like the majority of species – and the function of these structures – are undocumented. In short, why peristomes are so variable, and whether the various forms relate to prey capture, remains unknown.

**Fig. 1.**
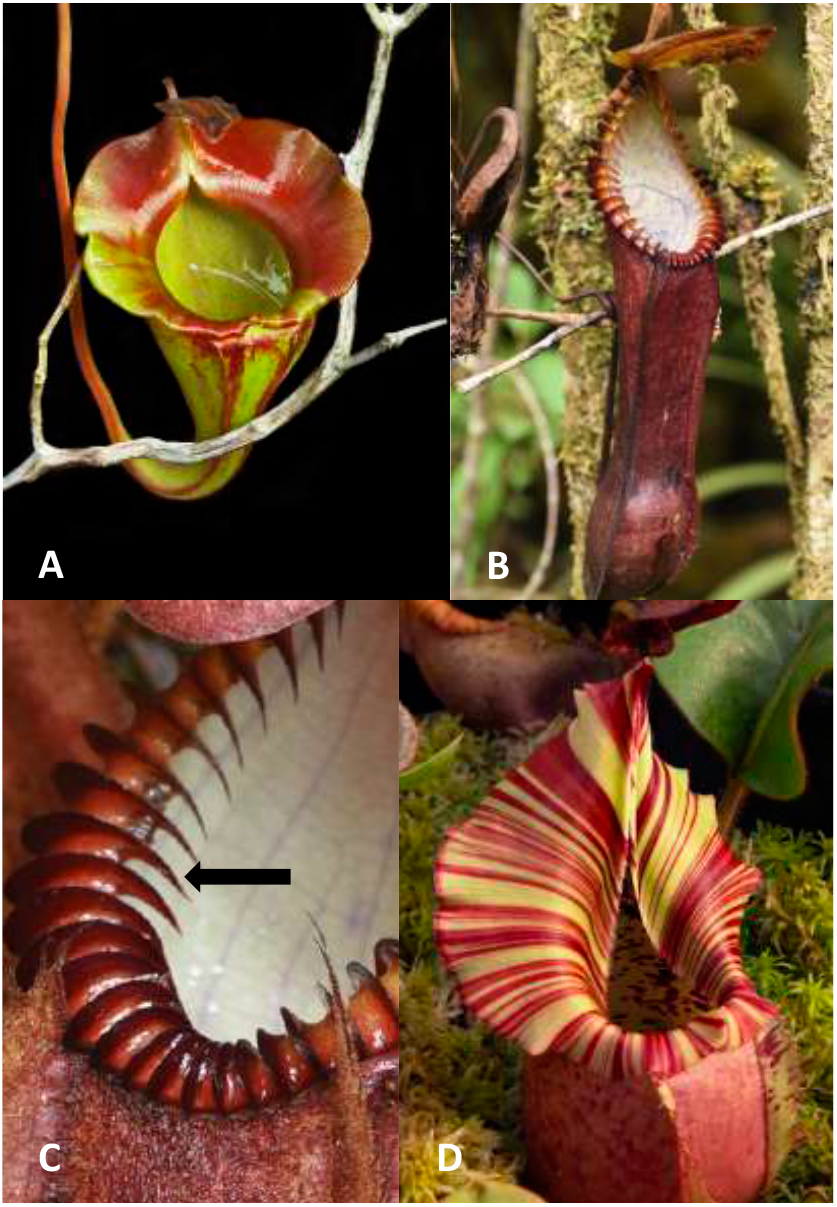
Divergent morphology in the genus *Nepenthes* shown by A, the flat peristome of *N. jacquelineae*; B-C the prominent teeth (arrow) of *N. hamata*, and the conspicuously flared peristome of *N. veitchii*. Photos A and D by Domonick Gravine; photos B and C by Jeremiah Harris.

**Fig. 2.**
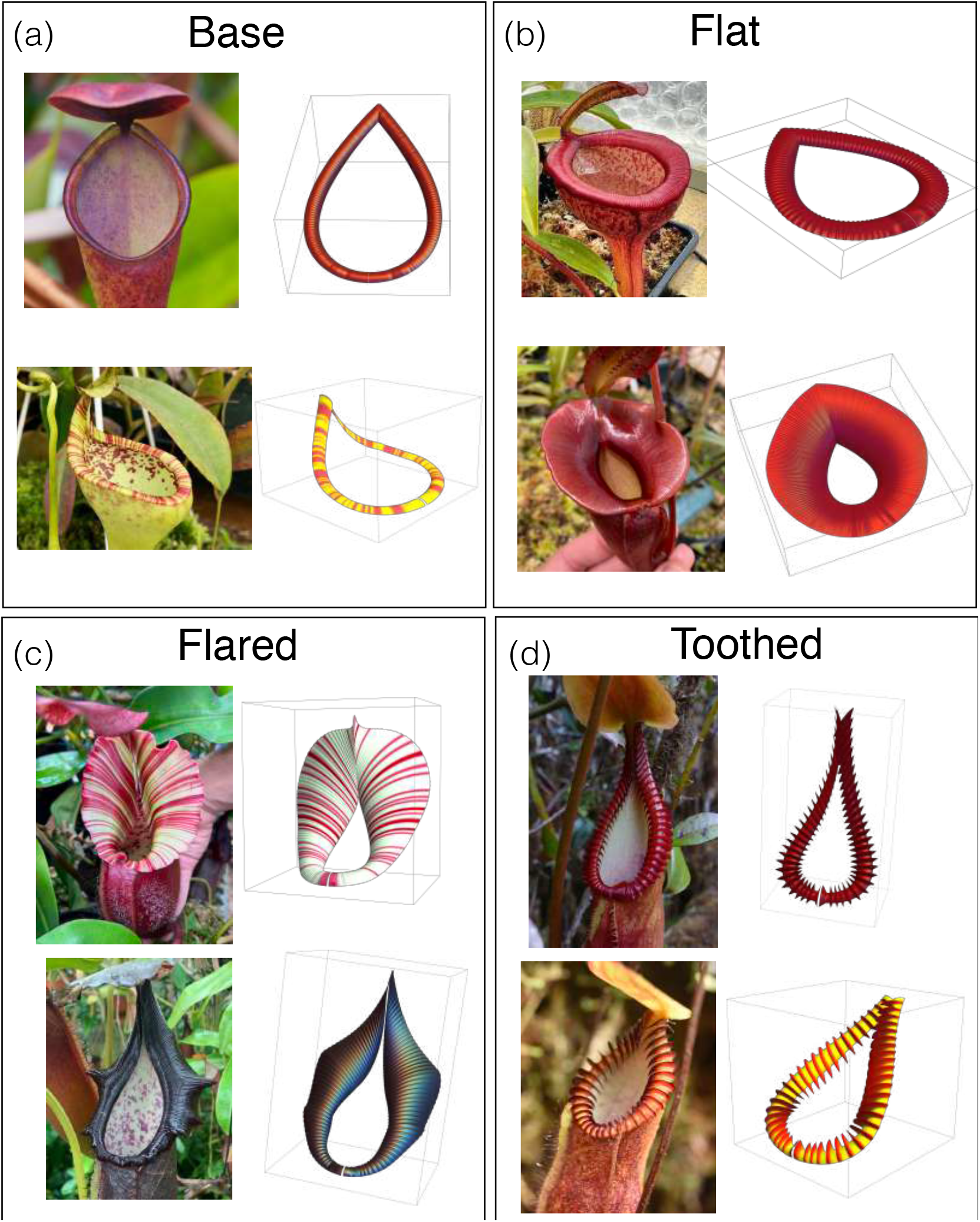
Variation in peristome geometry and mathematical reconstructions. We categorize peristomes into 4 categories: (a) Base geometry, exemplified by *N. pervillei* (top) and *N. eymae* (bottom); (b) Flat geometry, exemplified by *N. jamban* (top) and *N. jacquelineae* (bottom); (c) Flared geometry, exemplified by *N. veitchii* (top) and *N. naga* (bottom); (d) Toothed geometry, exemplified by *N. macrophylla* (top) and *N. diabolica* (bottom). Mathematical surface reconstructions for each peristome are shown at right. Details on the mathematical construction process given in SM Section 1. *N. pervillei* photo by Ulrike Bauer; *N. eymae* photo by Sarracenia Northwest; *N. jamban, N. naga*, and *N. macrophylla* photos by Tom Bennet (tomscarnivores.com); *N. jacquelinae, N. veitchii*, and *N. diabolica* photos by Jeremiah Harris.

Here we present the first mathematical framework to link divergent three-dimensional peristome geometries to the physics of prey capture. Linking form and function, we test the hypothesis that shape and ornamentation, orientation in a gravity field, presence of teeth, and peristome size, influence the diversity of prey capture in *Nepenthes*.

## 1. Mathematical approach

Our objective is to develop and analyze a mathematical framework linking peristome geometry to prey capture to investigate whether the observed diversity in peristome geometry can be understood in simple physical terms relating to prey-capture functionality. Of the 179 known species, there exists a wide variety in peristome size and morphology. Here we focus on three key geometric features of the peristome: i) the presence and degree of peristome *flaring* broad and often fluted, ii) the *orientation*, or tilt, of the peristome with respect to gravity, and iii) the presence of surface features such as *ribbing* or in extreme cases, *teeth* prominent spine-like, parallel features. Based on these divergent features, we classified *Nepenthes* peristomes into four categories that could be easily compared, as illustrated in Fig. 2: Base, Flared, Flat, and Toothed. The Base geometry has a thin peristome, a roughly 45^*?*^ tilt with respect to the vertical, and inconspicuous ribbing. This type is exemplified by *N. pervillei*, a species from the Seychelles, established to be sister to all other species of *Nepenthes* (15). It is reasonable to assume that other, more ornate patterns of geometry, evolved from this ancestral state. Flared peristomes are similar to the base geometry distally (at the front), but flare out to varying degrees proximally (near the point of attachment to the lid). Flat peristomes have a similar geometry to Base, but with a wider rim; these are distinct from the flared peristomes in that the peristome is more uniform in width. These peristomes are also characterized by a flatter orientation with respect to gravity compared with the other types which are tilted such that the proximal region is lower than the distal portion. Toothed peristomes also have a similar geometry to Base – thin and without flaring – but possess prominent ribs, so large that they are often referred to as ‘teeth’, protruding outward from the peristome and projecting into the pitcher interior. Despite their conspicuousness, their function is unknown.

To fully explore the potential functions of the suite of features described above, we must first establish a robust mathematical framework that can accurately describe the diverse geometries involved. In Supplementary Material (SM) Section 1, we have outlined a systematic procedure for creating explicit, parameterized mathematical surfaces that model various peristomes. This approach allows us to efficiently generate realistic peristome geometries that can be modified easily and continuously as needed. Sample examples of these surfaces can be found in Fig. 2. By employing this construction process, we can create a wide range of peristome shapes and configurations and investigate their properties and functions. For a given peristome type, we have a vector of parameters *S* that defines the peristome surface ∑ *⊂* ℝ ^3^.

Given a peristome surface ∑(𝒮), we characterize prey capture capabilities by first considering the sliding of a point mass on ∑ as a function of surface wetness. Since we can neglect the deformation of the peristome due to the small mass of the insect, we assume that the peristome remains fixed and rigid. The first question is: *is an insect’s position* **p** ∈∑ *on the surface stable under the force of gravity*? This is a simple geometry problem that involves determining the local peristome orientation in the gravitational field using the normal vector **n** to ∑, and the coefficient of static friction.

The effect of increasing wetness is to reduce the stability of most positions. Therefore, our second question is crucial: *if a position on the peristome is unstable, will the insect slide into or out of the pitcher?* The dynamics of a point mass on the peristome is given by a system of differential equations that can be integrated in time until either the inner or outer edge of the peristome is reached. Points whose trajectory leads to the inside rim of the peristome will be deemed caught by the pitcher, contributing nutrients to the plant, while points whose trajectory leads to the outside rim will fall off the edge, contributing nothing.

Details outlining this procedure and our computational approach can be found in SM Section 2. Our explicit surface parameterization enables us to efficiently and straightforwardly calculate surface stability and sliding dynamics. Based on these computations, we can divide the surface ∑, for a given friction coefficient, into different non-intersecting regions of total area 𝒜 = 𝒜_stable_ + 𝒜_unstable_ = 𝒜_stable_ + 𝒜_in_ + 𝒜_out_ and:

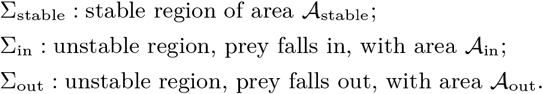

Next, we use the above approach to analyze flaring, orientation, and ribbing features. It is important to highlight the modeling trade-off: the analysis in these sections is carried out on detailed and realistic geometries, but using a highly idealized and simplified description of the insect itself as a point mass. To complement this analysis, we will present a second model to take into account the effect of *prey size* on capture capabilities.

## 2. The benefits of a flared peristome

The pitcher plant species *Nepenthes veitchii* is known for its striking peristome, which is broad and oblique. This peristome type is also observed in other species of *Nepenthes*, including *N. nebularum, N. hurrelliana, N. naga*, and *N. robcantleyei*. However, the prey spectra of these species in their natural habitats remain undocumented, and the evolutionary drivers behind this peristome morphology are still unknown.

To gain insight into the potential benefits of a Flared peristome for prey capture, we first analyze the stability properties of the peristome surface as wetness increases. By examining the peristome geometry and its response to different levels of wetness, we can develop a better understanding of how this structure functions and how it may have evolved to suit the needs of the plant.

In Fig. 3(a), we present the result for our model of a Flared peristome, with each point of the surface colored according to the vantage of the insect giving both its stability and dynamic properties: points in the region ∑_stable_ are green (safe); points in ∑_in_ are labeled red (unsafe), and points in ∑_out_ are labeled black. The different surfaces correspond to differing degrees of ‘slipperiness’: the friction coefficient, denoted *µ*, decreases following the arrow, corresponding to a more slippery surface.

**Fig. 3.**
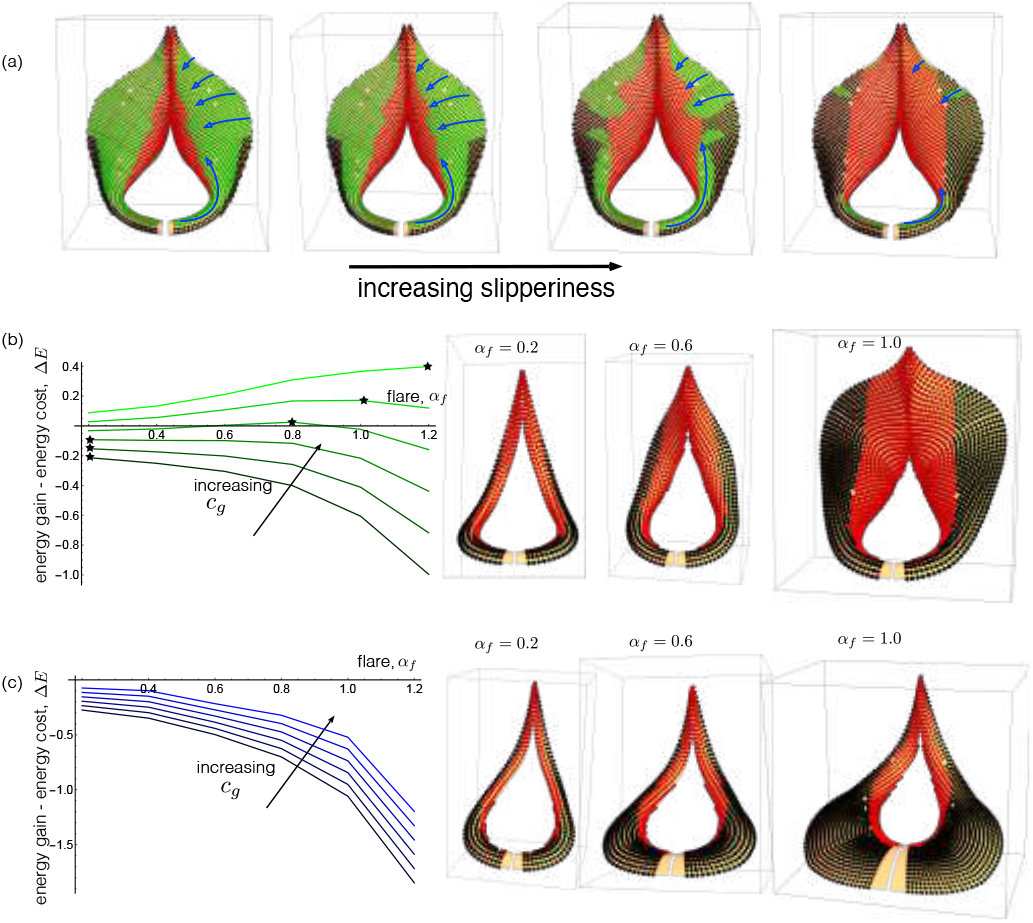
The impact of flaring on prey-capture. (a) Stability and capture properties of a Flared peristome as friction coefficient *µ* is decreased. Green points are stable, red points slide into the pitcher, and black points slide out. Stability ‘corridors’ – stable paths from the edge of the peristome to the unstable inner rim – are highlighted with blue arrows. (b), (c): Net energy gain Δ*E* plotted for increasing degree of peristome flaring, *α*_*f*_, and for different values of energy benefit parameter *c*_*g*_, for normal flaring (b) and lower rim flaring (c). The point of maximum Δ*E* is denoted with a star. Right: the peristome geometry at indicated values of *α*, with fall-in and fall-out points shown in red and black, respectively.

Naturally, as the surface becomes more slippery, a larger area becomes unstable; indeed ∑_stable_ shrinks to a set of zero area in the limit of zero friction. It is also unsurprising that points on the inner rim, where the surface becomes nearly vertical, are red (the dynamics end with the prey falling in), with this red region ∑_in_ expanding with increasing slipperiness. The black region, ∑_out_, is “useless” to both the plant and insect, as prey located at these points will fall out of the pitcher. It is interesting to note that ∑_out_ remains relatively small until very high slipperiness, and always has smaller total area than ∑_in_.

Nectar glands are located near the inner rim region of the peristome. Therefore it is in this general direction that preys are likely to be attracted. Further, a recent study (16) presents a capture mechanism in which scout ants are able to walk on the peristome surface without sliding and falling in; these scout ants recruit workers, enabling a batch catch and thus greater benefit than if the scout ant had fallen in. In the context of these two points, Flared geometry may be adaptive for capturing walking prey such as ants. At low slipperiness, there are few black regions; thus the surface geometry provides a safe platform for scout ants to locate nectar, and subsequent worker ants to follow pheromone trails to the red region. As slipperiness increases, stable green ‘corridors’ enable insects to walk from the outer edge of the peristome to the red region, as highlighted by blue arrows in Fig. 3(a). Owing to the climbing habit of *Nepenthes veitchii*, the proximal portion of the peristome often touches the vertical axis of the supporting tree. Here the flared peristome may act as a corridor to the pitfall trap – a form of shuttle for insects crawling up and down the tree.

### Energy considerations

A fundamental trade-off exists in carnivorous plants: leaves are modified into traps at the expense of photosynthetic efficiency because the traits of an effective insect trap are incompatible with those of an efficient light trap (4). Our analysis of Flared peristomes indicates a trade-off between prey capture and production. The peristome contributes little to photosynthesis, and is costly to construct (17), suggesting a strong selective advantage to such a structure in an environment that is nutrient-stressed in the first place. Quantifying such trade-offs between peristome investment and prey capture with empirical data is challenging, not least since the identity of prey in nature is unknown for most species. However, we can nevertheless gain insight into this problem by usin a modeling approach in which we assume that the energetic benefit, denoted *E*_gain_, is an increasing function of the capture surface area; that is

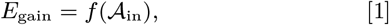

where *f* is a monotonically increasing function. This models the assumption that the benefit increases with the number of prey caught, and that the number of prey caught increases with the area of peristome from which prey fall. Since 𝒜_in_ depends on the friction coefficient, to simplify our analysis we compute 𝒜_in_ in the case of a perfectly wetted surface (*µ* = 0). We model the energetic cost as an increasing function of the total peristome area, that is

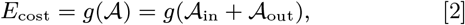

the latter equality reflecting the fact that the stable area shrinks to zero in the limit of *µ* → 0.

We can then define the net energy

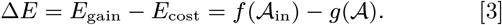

We want to express Δ*E* as a function of a given peristome feature that may be varied through natural developmental mechanisms. Then, evolution through natural selection should serve to vary this feature to the point where Δ*E* is maximal. If changing a given feature decreases Δ*E*, we expect to see such changes in nature. Of course it will depend on the specific form of the functions *f* and *g*. Here we consider a generic form 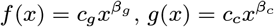, where the constants *c*_*g*_ and *c*_*c*_ characterize the energetic impact of increased capture area and total area, respectively, while the exponents *β*_*g*_ and *β*_*c*_ characterize possible non-linearity in the pathway between areas and energy.

We now examine flaring under this framework. Our construction method enables to continuously vary the degree of flaring, from thin (as in Base) to a widely flared peristome, or even to that beyond what is observed in nature. Therefore, we express Δ*E* as a continuous function of the flaring parameter *α*_*f*_ where *α*_*f*_ ranges from 0.2 (unflared) to 1.0 (typical flaring observed in *N. veitchii*) (details in SM Section 1). For a given *α*_*f*_, we seed the peristome with a uniform distribution of point masses, integrate forward the dynamic trajectories, and compute the capture (and miss) areas as fractions of total area based on the number of trajectories leading to the inner (and outer) rim (details in SM Section 2). In Fig. 3(b), we plot Δ*E* over a range of values of *α*_*f*_ for varying choices of *c*_*g*_, where we have fixed without loss of generality *c*_*c*_ = 1, and with other parameters taken to be *β*_*c*_ = 1, *β*_*g*_ = 1.1. For each choice of *α*_*f*_ the maximum of Δ*E* is denoted with a star. For low values of *c*_*g*_, Δ*E* decreases monotonically with *α*_*f*_. Here the benefit from increased prey capture is relatively low: the cost of increased total area outweighs the gain from increased capture area; for a species with these parameters, it would not be energetically favorable to increase flaring. For an increased *c*_*g*_, however, Δ*E* exhibits non-monotonic behavior, and indeed with an interior maximum, the degree of flaring to which our model would predict selection pressures that will drive the selection of this feature.

One great advantage of modeling is that it allows us to investigate features that are not found in nature. For instance, in Fig. 3(c), we repeated the same analysis, but with flaring along the bottom rim of the peristome. Such peristome geometries are not observed in nature, and our energy model demonstrates why this might be the case: the increased area at the bottom rim does not contribute to prey capture, as prey located there will fall out of the pitcher when slippery. Thus, increasing flaring in this manner does not result in a net benefit. This is evidenced by the fact that Δ*E* decreases as flaring increases for all the tested values of *c*_*g*_, rendering it a non-adaptive feature.

## 3. Peristome orientation

Next we consider the orientation of the peristome with respect to the vertical. Peristome orientation varies conspicuously across the genus from near-horizontal, for example in *N. jamban* and *N. jacquelinae* [Fig. 2(b)], to an orientation of c. 45^*?*^, for example *N. veitchii* and *N. truncata* [Fig. 2(c)].

To determine the relevance of peristome orientation to prey capture, we have varied this angle, defined as *ϕ* in our construction (see SM Section 1), from being flat (*ϕ* = 0) to vertical (*ϕ* = *π/*2 = 90^*?*^), while also varying the friction coefficient *µ*. Considering again the Flared peristome model, Fig. 4(a) shows how the regions ∑_stable_ (green), ∑_in_ (red), and ∑_out_ (black) vary both with tilt and friction coefficient. Comparing the top and bottom rows, it is evident that tilt has a strong impact on stability in the case of flat or vertical peristomes, and such that very little is changed by varying *µ*. This is in sharp contrast to the Base middle case *ϕ* = *π/*4 = 45^*?*^. In the context of stability corridors towards the unstable red zone (Section 2) that disappear as wetting increases, this trend suggests that this strategy will be most successful at an intermediate tilt.

**Fig. 4.**
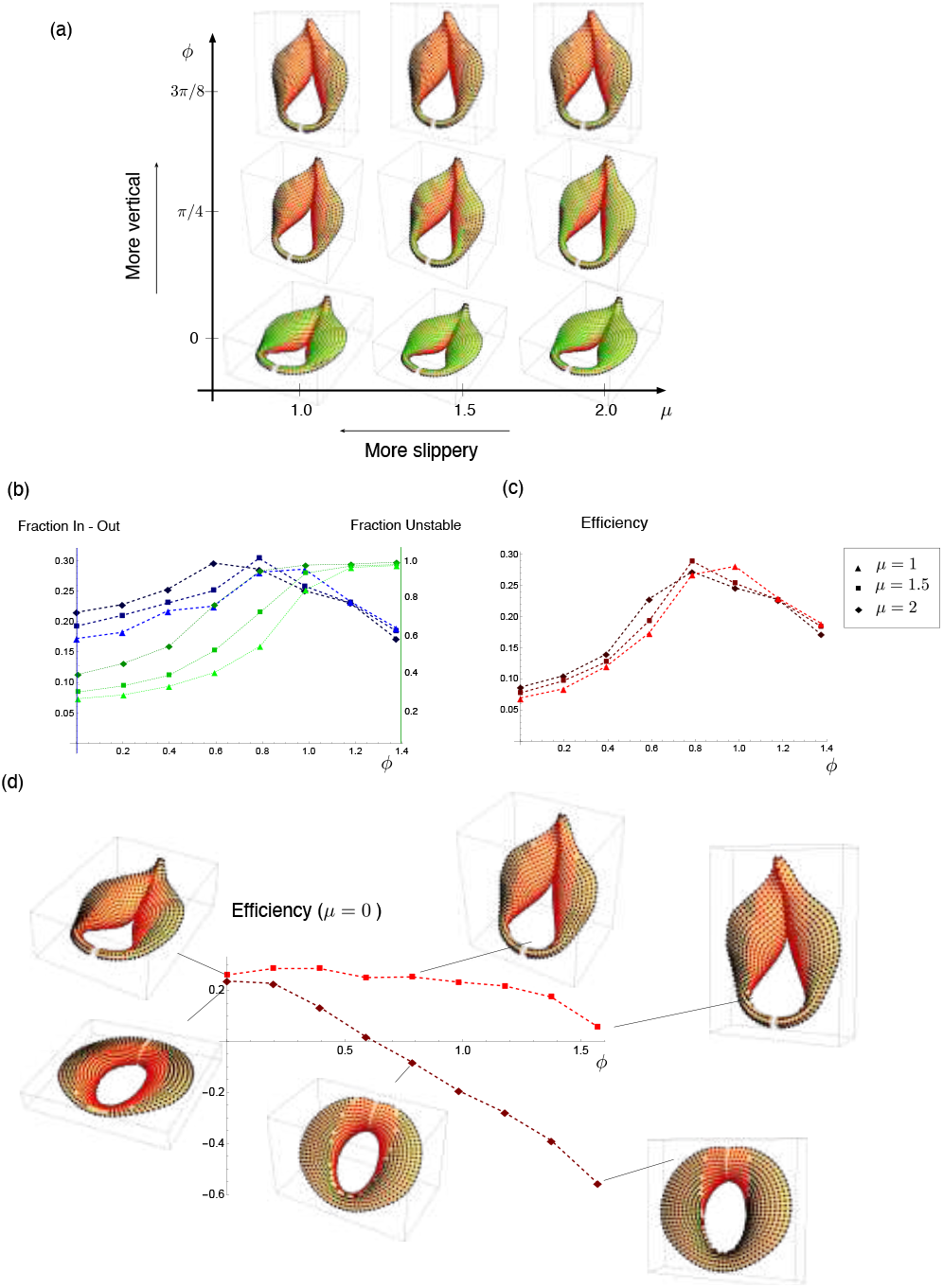
The impact of peristome orientation on prey-capture. (a) A phase diagram showing stability and capture properties for varying friction coefficient *µ* and peristome tilt with respect to the vertical, *ϕ*, for a model of a flared peristome. Green points are stable, red points slide into the pitcher, and black points slide out. (b) Plots of ℱ unstable (green) and ℱ in-out (blue) as a function of tilt *ϕ* for the flared peristome, each for varying values of *µ*, as indicated. (c) Capture efficiency measure as a function of *ϕ* and varying values of *µ*. (d) Efficiency measure as a function of tilt for a fully wetted peristome (*µ* = 0) for the flared peristome model (bright red) curve and a model of *N. jacquelineae* (dark red) displying a less flared and more uniform peristome geometry. Red points slide into the pitcher and black points slide out.

To quantify the benefit of a given orientation, we define the following metrics:

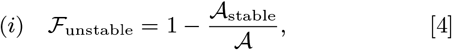

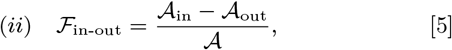

where ℱ_unstable_ is the fraction of the surface that is unstable, while ℱ_in-out_ is the difference between the fraction of the surface that is unstable and for which dynamic motion leads to falling in and the unstable fraction for which dynamics leads to falling out. These are computed for the Flared peristome in Fig. 4(b), with ℱ_unstable_ plotted as the green lines and ℱ_in-out_ appearing as blue lines, each for 3 different values of *µ*. The unstable fraction increases monotonically, such that almost the entire surface is unstable at the vertical orientation *ϕ* = *π/*2, while ℱ_in-out_ shows a non-monotonic relation with tilt.

From these metrics, we then compute an efficiency

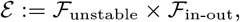

defined as the product of unstable fraction and ‘in minus out’ fraction. A surface with perfect efficiency with = 1 is such that every point falls in. Note that with this definition, negative efficiency is possible when more points fall out than in. The efficiency metric is plotted in Fig. 4(c), and interestingly we see has a maximum value near *ϕ* = *π/*4, i.e. the 45^*?*^ tilt that is observed in nature, for all values of the friction coefficient.

However, as noted above, not all species exhibit a 45^*?*^ tilt. For instance, the peristome of *N. jacquelineae* is oriented much closer to the horizontal (*ϕ* = 0 in our description). The peristome of *N. jacquelineae* is also distinctly different from that of *N. veitchii*, with a flatter and more uniform shape, and only a slight gradient towards the center. In Fig. 4(d), we plot the efficiency metric against tilt for our model of *N. jacquelineae*. Since the peristome is flat, the surface must presumably become very slippery for any points to become unstable; for this calculation, then, we have thus set the friction coefficient to zero, so that all points on the surface (except for a curve of zero area) are unstable. For reference, we also include the same calculation for our model of the Flared peristome. Plotted on this scale, and for a completely slippery surface, the efficiency is nearly constant for *N. veitchii*, showing only a noticeable decrease at the highest tilt. The efficiency of *N. jacquelineae*, on the other hand, decreases significantly and monotonically with increasing tilt, reaching negative values before a 45^*?*^ tilt and with nearly 60% more points falling out than in at vertical. Because the peristome shape is flat, it requires significant wetting to capture any prey, but then the slight gradient in the geometry is best suited for capture with zero tilt; as the peristome tilt increases, more and more points slide off the bottom instead of to the inside.

Our model thus predicts a strong link between tilt and prey capture, but in a non-trivial way, with the optimal tilt itself a function of the peristome shape. Taken together, this indicates that tilting may be an adaptation to optimize prey capture efficiency.

## 4. On ribs and teeth

All peristome surfaces possess ribs of varying height and wavelength. In a handful of species these ribs are highly conspicuous and blade-like (referred to as teeth), for example in *N. macrophylla, N. diabolica* [Fig. 2(d)], and *N. villosa* and *N. hamata*, (not shown). Phylogenomic data indicate this phenomenon has evolved independently in the genus *Nepenthes* (15). In this section, we examine the prey-capture benefit that may be obtained from such features, in the context of a cost-benefit analysis. Typically, ribs have sharp peaks and wider smooth valleys. Intuitively, the presence of ribs is beneficial as prey that may have slid off the external pitcher are instead guided into the trap. However, such features increase the area at a significant energetic cost. Following Section 2, we quantify the energetic cost and benefit trade-off using Eqs. (1) and (2) to define the energetic gain *E*_gain_ in terms of capture area and energetic cost *E*_cost_ in terms of total surface area. As before, the metric of relevance is the net energy Δ*E* = *E*_gain_ −*E*_cost_ [Eq. (3)]. Here we examine Δ*E* as a function of a single parameter characterizing the size of the teeth (the wavelength is fixed based on observations of living material). We first consider the presence of ribs within a Flared peristome. In Fig. 5(a), we vary the rib height, denoted ϵ, from ϵ = 0 (perfectly smooth) to ϵ = 0.75, corresponding to a rib height greater than that observed in samples of *N. veitchii*. We have used the same form of energy functions *f* and *g* as in Fig. 3(b), and have fixed *c*_*g*_ = 2, which is the value at which Δ*E* attained a maximum at the flaring value *α*_*f*_ = 1.0. For these values, Δ*E* reaches a maximum at a ribbing height similar to that observed in nature. Clearly, there is a limit that is reached when the construction cost of increased rib height outweighs the benefit of prey capture.

**Fig. 5.**
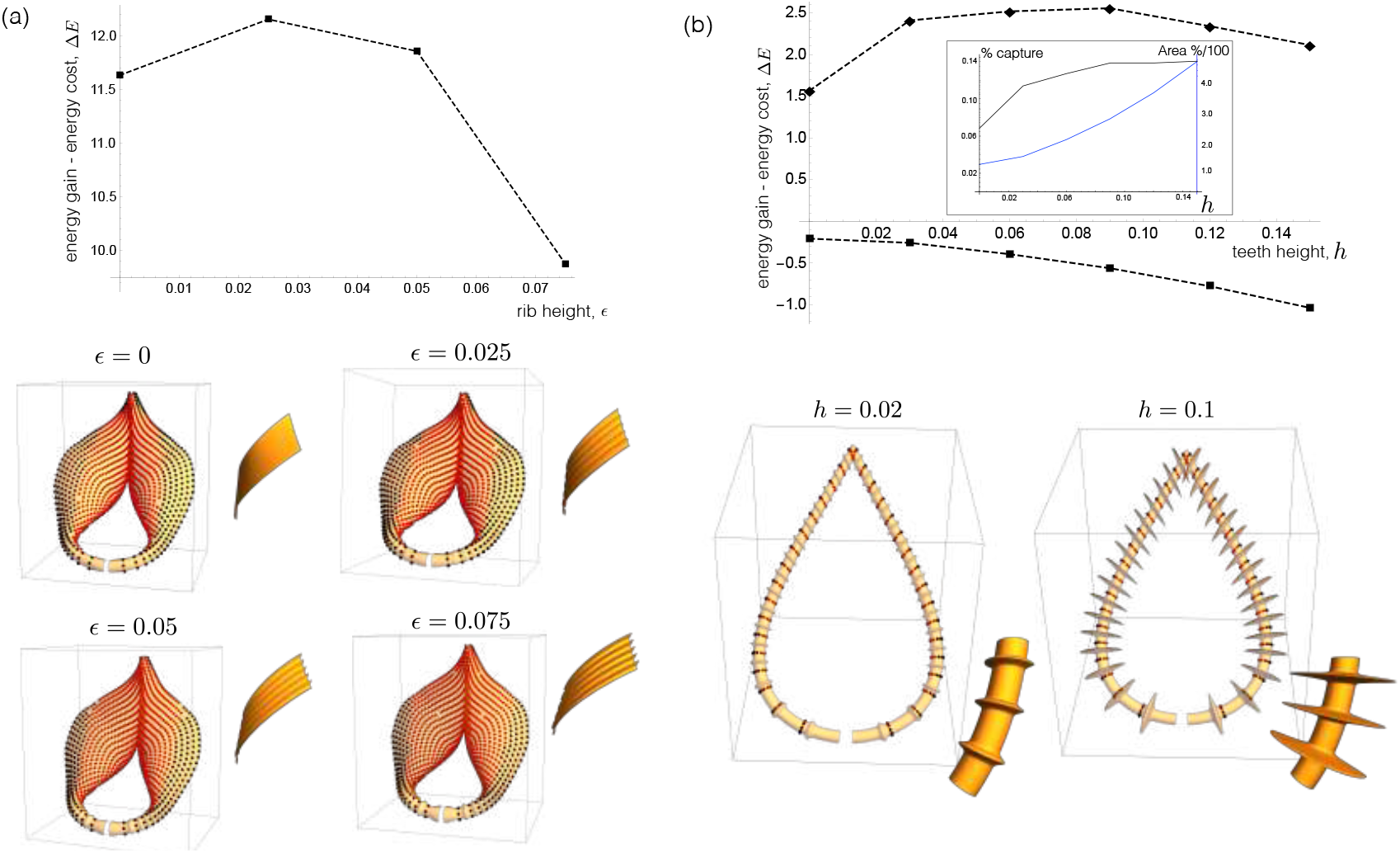
Impact of ribbing and teeth features on prey capture. (a) Net energy Δ*E* plotted against rib height for a model of a flared peristome. Right: capture properties for a perfectly wetted surface (*µ* = 0); red points slide into the pitcher, and black points slide out. Zoom in of surface ribbing shown for each height. Energy parameters are *c*_*g*_ = 1, *β*_*g*_ = 1.1, *β*_*c*_ = *c*_*c*_ = 1. (b) Net energy Δ*E* plotted against teeth height for a model of *N. hamata*. The lower graph, with square markers, has the same energy parameters as in (a); the upper graph, with diamond markers, has hugely increased energy gain, *c*_*g*_ = 50, *β*_*g*_ = 0.8, other parameters equal. Right: capture properties for a perfectly wetted surface (*µ* = 0); and zoom in to show teeth features at the indicated heights.

Comparing Fig. 3(b) and Fig. 5(a), we emphasize that using the same energy functions with equivalent constants, our model predicts optimal levels of flaring and ribbing that are consistent with those observed in nature. This adds weight to the hypothesis that these features confer a selective advantage in the capture versus construction trade-off.

In Fig. 5(b), we perform the same analysis for a model of a thin peristome with varying heights of teeth *h*; in the case of large teeth, these are based on *N. hamata*. The net energy Δ*E* is plotted against *h* for the same parameter values as above, appearing as the dashed curve with square markers. For these values, Δ*E* decreases monotonically with *h* and there is no net energetic benefit associated with producing teeth. The inset plots both the fraction of seeded points captured (black) and the surface area divided by the initial area (blue). While teeth do increase the capture fraction, it only does so by a small margin, while the area increases by a factor of 4 at the greatest height of teeth. In other words, the cost significantly outweighs the benefit. Since the construction cost is considerable, it is possible that teeth serve a function that falls outside the scope of our model, for instance, retention of prey. The ends of the teeth project markedly into the interior pitcher and could form a barricade that could prevent large prey from escaping. We should note that the presence of such a prominent feature can be predicted in our framework, but only if the energetic gain of any increased capture is weighted highly. For instance, the dashed line with diamond markers in Fig. 5(b) plots Δ*E* with *c*_*g*_ increased from 2 to 50, and *β*_*g*_ decreased to 0.8. Here an interior maximum at a realistic teeth height for observed species is attained, though we stress a 25-fold increase was required in the energetic gain parameter *c*_*g*_.

## 5. On peristome size

Finally, we explore the effect of peristome size on the efficiency of prey capture. Peristome dimensions vary across the genus, which could be a consequence of divergent selective pressures from differences in prey size and availability.

The point-mass model is scale free. Thus, in order to investigate the specific effect of prey size, we consider a minimal representation of a prey with finite size, sitting on a crosssection of a peristome. The peristome is modeled as a circle in a vertical plane, with radius *R* ≡1, taken to be a reference length. The prey is modeled as a rigid body in contact with the peristome at two points located at the same distance *ρ* from the rigid body’s center of mass *G*, and with angle 2*α* between *G* and the two contact points [Fig. 6(a)]. The scaled length *ρ* defines the lengthscale of the prey, while *α* characterizes its shape (long insect have larger *α* than compact ones). The position of the prey on the peristome is given by *θ*∈ [0, *π/*2], the angle between the vertical axis and the prey axis. We assume that the prey is only subject to its own weight *mg*, applied at the center of mass *G*. As before, we consider dry friction between the prey and the peristome, with coefficient *µ* at both contact points, and we derive the critical angle for frictional stability (see Refs. 18, 19, and SM Section 3). In the case of a prey with finite spatial extent, another instability may occur where the prey loses contact with the surface and tumbles without slipping into the trap, which will occur when *θ > α* [Fig. 6(b)]. For each value of *ρ* and *α*, and for a fixed friction coefficient *µ*, we compute exactly the maximum angle *θ* = *θ*_*c*_ beyond which equilibrium is lost, and the prey either slips or tumbles. The result is plotted in Fig. 6(c), where *θ*_*c*_ appears as a color map in the *α*-*ρ* plane – here, blue corresponds to *θ*_*c*_ = 0, i.e. vanishing stable zone, while yellow corresponds to the largest stable zone (with *θ*_*c*,max_ = arctan *µ*, corresponding to the point-mass limit). A region in which tumbling occurs before slipping is indicated on the left side of the plot, for small *α*. The uncolored white region corresponds to disregarded points in which the leg axis would have to penetrate the surface.

**Fig. 6.**
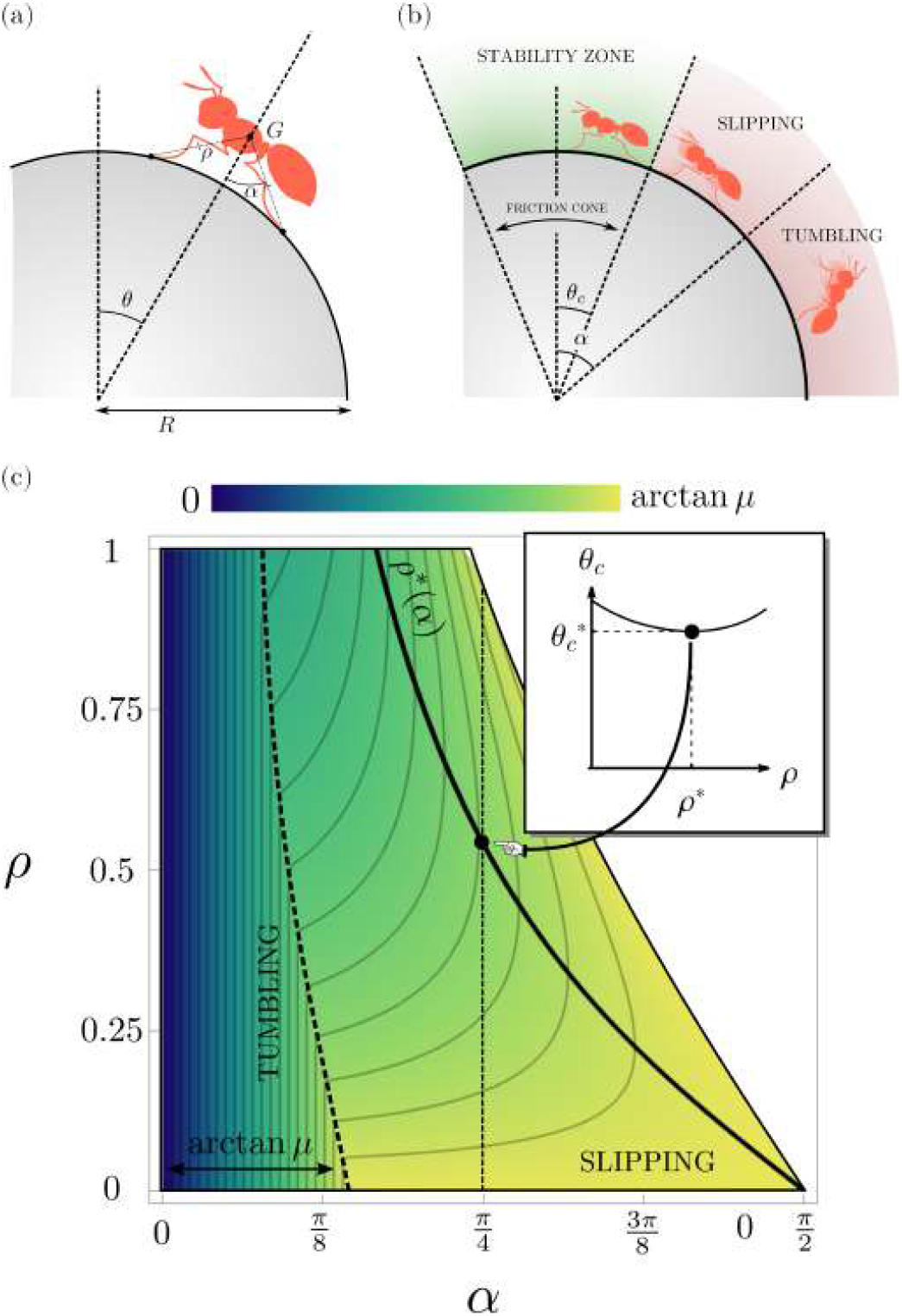
Finite-size prey. (a) Schematic of the two-leg prey model geometry. (b) The two modes of capture: slipping and tumbling. The friction cone with angle *θ*_*c*_ characterizes the zone where frictional stability can be maintained for a given prey and friction coefficient *µ*. (c) Density plot showing the size of the stability zone (*θ*_*c*_) vs prey angle (*α*) and prey size (*ρ*), with *µ* = 0.5. Inset: plot of *θ*_*c*_ vs *ρ*. for *α* = *π/*4. Note that 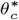 (*ρ*) has a minimum *θ*^*c*\*^, reached at a finite value *ρ*^*^.

From the perspective of a prey, Fig. 6(c) shows that it is advantageous to be as flat as possible, in the sense that for any *ρ*, the largest stability angle is achieved when *α* is maximal. It is also generally the case that small preys have an advantage. For any *α*, the stability zone *θ*_*c*_ is maximal when *ρ*→ 0, which shows that the point-mass model provides a lower bound for the trapping efficiency. However, as *ρ* increases there is a nonlinear relation between size and stability (in the slipping regime). This is evident in the inset of Fig. 6(c), which plots *θ*_*c*_ against *ρ* for *α* = *π/*4. Here *θ*_*c*_ is non-monotonic, achieving a minimum value at an intermediate size, denoted *ρ*^*^ (an exact expression is provided in SM Section 3). This reflects the notion that larger legs may reach further around the peristome, with stability increasing up to the point where the legs are tangent to the surface.

Since we have scaled the insect length by the peristome size, for a given insect shape given by *ρ* and 0 *< α < π/*2, the insect lengthscale is *r* = *ρR*. Therefore, there is an optimal peristome size *R*^*^(*α*) = *r/ρ*^*^(*α*) that will be most effective at capturing the prey. Note that, in the slipping regime, the optimum size *ρ*^*^ is independent of the frictional properties of the peristome and is therefore a universal geometric property of the model.

For instance, considering an environment where typical preys have angle *α* = *π/*4 and size *R*, and arbitrary but small friction coefficient *µ≪* 1 (slippery peristome), we have *ρ*^*^≈0.5, and the highest trapping efficiency will be achieved by peristomes with *R* ≈2*ℓ*, which generates a 17% efficiency gain with respect to the most stable case *ρ* →0, all other things being equal. From an evolutionary viewpoint, this observation suggests the existence of a linear scaling law between the peristome size and the typical size of the preys that will be most easily caught in a particular ecological niche. A few studies (20, 21) have classified prey contents for a range of *Nepenthes* species in a given habitat, and these seem to be consistent with a correlation between larger peristomes and larger prey, e.g. pitchers with small peristomes, on the order of *R* ≈1 mm in (*N. albomarginata* and *N. gracilis*) almost exclusively captured termites and ants, while pitchers with larger peristomes, on the order of *R*≈ 5–10 mm or more (e.g. *N. gigantea, N. rafflesiana*, and *N. hemsleyana*) also captured ants, but also captured a wider variety of other prey, including Gasteropoda, Coleoptera, and Arachnida. However, these data do not include measurement of the actual size of the prey trapped and the trend is therefore only qualitative.

## 6. Discussion

The remarkable diversity of trap forms in the genus *Nepenthes* is emerging as an adaptive radiation analogous to better-known examples from the animal kingdom, such as the beaks of Darwin’s finches (3). However the drivers of the adaptive radiation in *Nepenthes* remain poorly known or unexamined in most species. By using mathematical modeling and the laws of Newtonian mechanics, our study has revealed that prey capture is influenced both by peristome shape and relative size. Therefore the diversity of peristomes in *Nepenthes* appears to have evolved in response to dietary needs, adding weight to the hypothesis that a divergence in trap form represents an adaptive radiation.

Carnivory evolved independently in five orders of flowering plants in response to nutrient stress. Advances in genome and transcriptome sequencing have revealed the repurposing of defense-related genes is an important trend in the evolution of plant carnivory (22). *Nepenthes* evolved within a clade that includes snap trap leaves in the genera *Dionaea* and *Aldrovanda*, in which a touch-sensing mechanism allows rapid closure; and flypaper trap leaves which move more slowly for example *Drosera*. Active mechanisms represent geometric and mechanical solutions adapted for specific prey situations; accordingly, a high diversity of trap configurations has evolved across the various niches occupied by carnivorous plants (23). In the case of *Nepenthes*, prey capture relies on insects being attracted to, and sliding off, the wet peristome. Attraction is achieved by nectar and coloration, while sliding is achieved both by the surface properties of the peristome and the peristome geometry. While the surface properties have been well-documented, here we provide the first study linking geometry and mechanics to prey capture. Just as in active traps, efficacy is underpinned by both geometry and mechanics.

An optimal geometry might be expected to exist to enable passive capture irrespective of insect type or size. However we find no such evidence of this; on the contrary, our analysis provides a clear context in which we may understand why peristome geometry in *Nepenthes* is divergent. We consider the value of a given peristome feature in terms of cost-benefit: the energetic cost of peristome construction against the energetic gains of prey capture. While cost benefit is dependent on biotic variables, we provide a hypothetical framework for investigating this balance. In the case of peristome flaring, our analysis points to a consistent means by which an evolutionary path from a narrow to a flared peristome might exist. Moreover, our analysis may also provide an explanation for an evolutionary divergence in peristome geometry. Indeed, a small change in the parameter *c*_*g*_, which characterizes the relative energetic gain of increased prey capture, has a strong impact on the optimal flaring, and for some values the unflared geometry is energetically optimal. As the energy pathways are likely to vary among species, so will the optimal degree of flaring, and in this context it is not surprising that not all species possess widely flared peristomes. A similar situation exists in the case of ribbing or teeth features which, though they generally serve to increase the prey capture functionality, are also energetically costly to produce. Considering peristome orientation with respect to gravity, our analysis provides a plausible physical explanation for the correlation between geometry and orientation, demonstrating that a wider and more uniform peristome has a better capture efficiency when oriented horizontally.

In these examples, capture success was linked to geometric complexities, and a detailed geometric description was needed for which we sacrificed prey complexity in the description. We also analysed finite-sized prey with multiple contact points on a simplified constant curvature surface restricted to 2D. Here again, the connection between geometry and prey specifics was evident and we identified a nonlinear relationship between prey geometry and capture efficiency. Taken together, this hints at a fine-tuning of peristome size to optimize prey capture likelihood for a given shape and size.

The two distinct forms of analysis we have presented each incorporate simplifications in different ways. Amalgamating the approaches, i.e. combining three-dimensional geometries with a detailed description of finite prey possessing multiple surface contact points, would be more powerful; though it poses a significant challenge to do so in a tractable manner. While our analysis focused on the functional benefits of peristome size and geometry, a functional geometry is useless without a developmental process capable of generating it, and a complementary direction of future research would be the morphogenesis of peristome.

The striking divergence of pitcher forms in *Nepenthes* suggests that they should attract different prey across their various habitats. An investigation into the prey spectra of seven Bornean species indeed revealed different combinations of trapped ants, flying insects, termites, and non-insect organisms (21). Prey capture is also known to shift with altitude. Many lowland species are attractive to ants, and possess waxy interior pitcher surfaces effective for capturing these insects (9, 24). By contrast, montane species, which tend to have viscoelastic pitcher fluids, are more effective at trapping flying prey (10, 25). Beetles appear to be the most abundant prey for *N. villosa*, a montane species with conspicuous teeth (20). Peristome teeth may play a role in the retention of bulky prey; however, data from other species with prominent teeth are lacking. Different combinations of pitcher surface and fluid properties probably correlate with peristome size and geometry. For example pitchers without waxy surfaces often produce larger and more inward-sloping peristomes (24). But despite its central role in capture, we know virtually nothing about how prey shifts with changes in peristome morphology. Further work would benefit from empirical and observation data on prey capture from across a range of pitcher and peristome forms in different habitats.

Our study provides a mathematical construct for quantitatively linking geometry to prey capture. Investigating this link empirically is a crucial next step. Of course, prey capture will also depend on variables beyond geometry, such as coloration and nectar production; furthermore, pitcher morphology usually varies with plant age (traps produced by young rosettes are distinct from those on mature vines). In principle, our conceptual approach can accommodate the inclusion of such features. This highlights the value of mathematical modeling as an iterative process that can both motivate and adapt to new empirical studies. In conclusion, this approach provides a platform for testing hypotheses on the evolution of nature’s green predators: some of the plant kingdom’s greatest enigmas.

## Data availability

Mathematica notebooks reproducing model output will be made available in a public depository upon manuscript acceptance.

## Acknowledgments

The support for AG by the *Engineering and Physical Sciences Research Council* of Great Britain under research grant EP/R020205/1 is gratefully acknowledged. The authors also thank Tom Strube, who was involved in early discussions related to this project. For the purpose of Open Access, the author will apply a CC BY public copyright licence to any Author Accepted Manuscript (AAM) version arising from this submission.

