## Supplementary Information for "Mechanics reveals the role of peristome geometry in prey capture in carnivorous pitcher plants (*Nepenthes*)"

**This PDF file includes:**

Figs. S1 to S3
SI References

### 1. Peristome surface construction

#### A. General procedure.

**Overview.** We first outline the procedure used to generate realistic parameterizations of three-dimensional peristome surfaces. This process is shown schematically in Fig. S1, and consists of the following steps:

1. Define a planar curve $(x(s), y(s))$ for the base shape for the peristome.

2. Define an angle of inclination $\phi$ from the horizontal, i.e. $\phi = 0$ corresponds to a perfectly flat peristome, while $\phi = \pi/2$ for a vertical orientation.

3. Define a function $h_2 = h_2(s)$ describing deviation of peristome base shape from planarity. This was used in constructing models of peristome for which the proximal end is raised relative to the rest of the peristome.

4. Prescribe the cross-sectional shape of the peristome as a 2D curve, defined in polar coordinates by a function $r(\theta)$, at a finite number of points $\{s_1, s_2, s_3, \ldots, s_n\}$.

5. To create a surface from the set of discrete cross-sectional shapes, we create interpolating functions in $s$ from the values of the shape parameters at each point $s_i$, therefore generating a surface parameterized by $(s, \theta)$.

6. Additional features such as ribbing, teeth, or color patterns may then be added on top.

**More details on Step 4.** Most of the fine tuning for surface construction appears in Step 4, creating the cross-sectional shapes. This step involves a number of sub-steps that we outline further here. First, we note that from the parameterization of the base shape, and given the tilt $\phi$, we define the space curve

$$\mathbf{R}(s) = (x(s), \cos\phi\, y(s), \sin\phi\, y(s)). \quad [1]$$

The tangent ($\mathbf{T}$), normal ($\mathbf{N}$) and binormal ($\mathbf{B}$) vectors to this curve may then be defined at each point $s$ as follows:

$$\alpha(s)\mathbf{T}(s) = (x'(s), \cos\phi\, y'(s), \sin\phi\, y'(s)) \quad [2]$$

$$\alpha(s)\mathbf{N}(s) = (y'(s), \cos\phi\, x'(s), \sin\phi\, x'(s)) \quad [3]$$

$$\mathbf{B}(s) = (0, -\sin\phi, \cos\phi), \quad [4]$$

where $\alpha(s)^2 := x'(s)^2 + y'(s)^2$.

In order to create easily manipulated cross-sectional shapes, we first restrict to a class of functions for the shape, such that the precise shape is determined by a small number of parameters that we can fix at each point $s_i$. For most of the peristome surfaces, we have used logarithmic spirals, i.e. in polar coordinates the curves $r(\theta) = r_0 e^{f\theta}$. The parameter $f$ is used to characterize flaring – $f \to 0$ corresponds to an arc of a circle, with radius $r_0$, while a larger value of $f$ produces a more flared curve (increasing radius as $\theta$ increases).*

The shape is initially defined in the **N**-**B** plane, from which we then define two rotations: we rotate the curve about the binormal direction by angle $\varphi$, and rotate the curve about the axis normal to the plane by $\Theta$. For example, if $\varphi = 0$ and $\Theta > 0$, the curve is placed in the **N**-**B** plane, rotated by angle $\Theta$ about the tangent **T**.

The domain of each curve is given by $\theta_1 < \theta < \theta_2$, where the $\theta_i$ are also chosen at each point $s_i$. Combining the above, at each point $s_i$ we define the following parameters:

$$\mathcal{S}_i = \{h_{2_i}, r_{0_i}, f_i, \varphi_i, \Theta_i, \theta_{1_i}, \theta_{2_i}\}. \qquad [5]$$

**More details on Step 5.** For given parameters $\mathcal{S}_i$, $i = 1, \ldots, n$, we create interpolating functions, transforming each of the 7 parameters in $\mathcal{S}_i$ into functions in $s$. For example, we generate the flaring function $f(s)$ as an interpolating function passing through each of $\{(s_1, f_1), (s_2, f_2), \ldots, (s_n, f_n)\}$. From these interpolating functions, we then define the peristome surface $\mathbf{P}(s, \theta)$ as follows:

$$\begin{aligned}\mathbf{P}(s,\theta) =& \mathbf{R}(s) \\ & - r_0(s)e^{f(s)\theta} \sin\varphi(s)\big(\cos\Theta(s)\cos\theta - \sin\Theta(s)\sin\theta\big)\mathbf{T}(s) \\ & + r_0(s)e^{f(s)\theta} \cos\varphi(s)\big(\cos\Theta(s)\cos\theta - \sin\Theta(s)\sin\theta\big)\mathbf{N}(s) \\ & + \Big[h_2(s) + r_0(s)e^{f(s)\theta}(\sin\Theta(s)\cos\theta + \cos\Theta(s)\sin\theta) - r_0(s)\exp(f(s)(\pi/2 - \Theta(s)))\Big]\mathbf{B}.\end{aligned} \qquad [6]$$

The final term in this expression shifts the curve so that the baseline curve $\mathbf{R}(s) + h_2\mathbf{B}$ is part of the final surface. We found that this choice produced smoother surfaces that were easier to manipulate to have desired features.

For ease of computation, we exploit the bilateral symmetry inherent in peristome geometry, so that we need only define and perform computations on one half (the right half, say) of the peristome, with all sliding properties assumed to be identical.† Surface plots in the main text show both halves, where the left half follows a mirror symmetry with the right half.

Via the process outlined above, having fixed the base shape and orientation, the surface is defined by the $7 \times n$ parameters $\mathcal{S}_i$, $i = 1, \ldots, n$. Typically we have used $n = 6$ points to balance efficiency while maintaining sufficient degrees of freedom to control surface geometry properties. With $n = 6$, the 42 parameters were varied using the *Manipulate* command in Mathematica (1), which provides a graphical user interface that rapidly updates the surface geometry while continuously varying any of the parameters. In this way, by monitoring the surface shape and comparing with images of actual *Nepenthes*, parameter sets for the $\mathcal{S}_i$ were constructed for each of the model peristome surfaces.

**Continuous flaring.** In generating Fig. 2 of the main text, we have continuously varied the peristome flaring. To do this, we first defined a flaring function $f(s)$, as well as all other surface parameters. We then incorporate a scaling factor $\beta$, so that the flaring function is given by $\beta f(s)$; ranging $\beta$ produces a naturally varying flare while maintaining other surface properties unchanged.

* Note that in our construction, the positive $x$-axis for each cross-section is aligned with the normal vector, which points inside the peristome, thus $\theta$ increases from inside edge to outside edge.

† Mirror symmetry with sliding would not be maintained if the peristome were rotated along the bilateral axis; however for the vast majority of *Nepenthes* that we have examined, there is negligible rotation of this type.

**On surface features.** The process outlined above produces a smooth surface. Adding surface features such as ribbing or teeth is straightforward, as this can be defined by a variation in the cross-sectional radius as a function of $s$. For instance, to add ridges with wavelength $\omega$ and amplitude $\epsilon$, the term $r_0(s)\mathrm{e}^{f(s)\theta}$ is replaced by $r_0(s)\mathrm{e}^{f(s)\theta} + \epsilon\cos(\omega\lambda(s))^m$ where $m$ is an integer characterizing the sharpness of the ridges ($m = 1$ for perfectly sinusoidal ridges, while a large value of $m$ produces ridges with sharp peaks and wide valleys). The function $\lambda(s)$ is the arc length of the baseline curve, defined by $\lambda'(s) = \alpha(s)$; this term is needed to maintain a constant wavelength, as the base curve is not defined (necessarily) to be an arc length parameterization. Both ridges and teeth may be defined in the same way; the main difference being that with teeth we use a much larger value of both $\epsilon$ and $m$, e.g. $m = 4$ for ridges and $m = 40$ for teeth. Note that in defining ridges and teeth in this way, they are aligned with the cross-sectional curves; therefore the function $\varphi(s)$ is used to rotate this alignment, for instance to be more oblique towards the proximal end, as we have observed in many species.

Coloration of surfaces in Fig. 1 of the main text was purely added for visual purposes in comparison with real specimens. Color patterns were produced using the *ColorFunction* environment in Mathematica.

**B. Particular surfaces.** Specific values of the parameters and functions used for the peristome models can be found in the deposited Mathematica notebooks, which include the construction of each of the models appearing in main text Fig. 2.

#### 2. Point-mass model

Here we outline the procedure for stability properties and dynamic motion of a point mass on the surfaces as constructed above.

**A. Seeding the surface.** While some properties, such as stability under dry friction, can be readily computed as a continuum property at each point on the surface, computation of the dynamics requires integrating the equations of motion forward in time, which must be done individually for any given point on the surface. Therefore, in order to approximate the relative areas of the surface for each category – stable, unstable with dynamics leading to falling into the pitcher, and unstable with dynamics leading to falling out of the pitcher – we first seed the surface with a large number of points. In order to space the seed points evenly across the surface, we first compute the metric tensor $\mathbf{G}$, which has components

$$G_{11} = \frac{\partial \mathbf{P}}{\partial s}\cdot\frac{\partial \mathbf{P}}{\partial s},\quad G_{12} = \frac{\partial \mathbf{P}}{\partial s}\cdot\frac{\partial \mathbf{P}}{\partial \theta},\quad G_{22} = \frac{\partial \mathbf{P}}{\partial \theta}\cdot\frac{\partial \mathbf{P}}{\partial \theta}. \qquad [7]$$

Noting that the line element $\mathrm{d}S$ of a curve on the surface satisfies $\mathrm{d}S^2 = G_{11}\mathrm{d}s^2 + 2G_{12}\mathrm{d}s\mathrm{d}\theta + G_{22}\mathrm{d}\theta^2$, our approach for seeding the surface is then as follows:

1. Fix a step size $\delta$
2. Fix $s = s_1$ (distal end).
3. Seed points along the cross-section:
   (i) Fix $\theta = \theta_1(s)$, and seed a point
   (ii) Increment $\theta \leftarrow \theta + \frac{\delta}{\sqrt{G_{22}(s,\theta)}}$, and seed a point
   (iii) Continue until $\theta_2$ is reached.
4. Increment $s \leftarrow s + \frac{\delta}{\sqrt{G_{11}(s,\theta)}}$
5. Return to step 3.
6. Repeat until $s \geq s_n$ is reached.

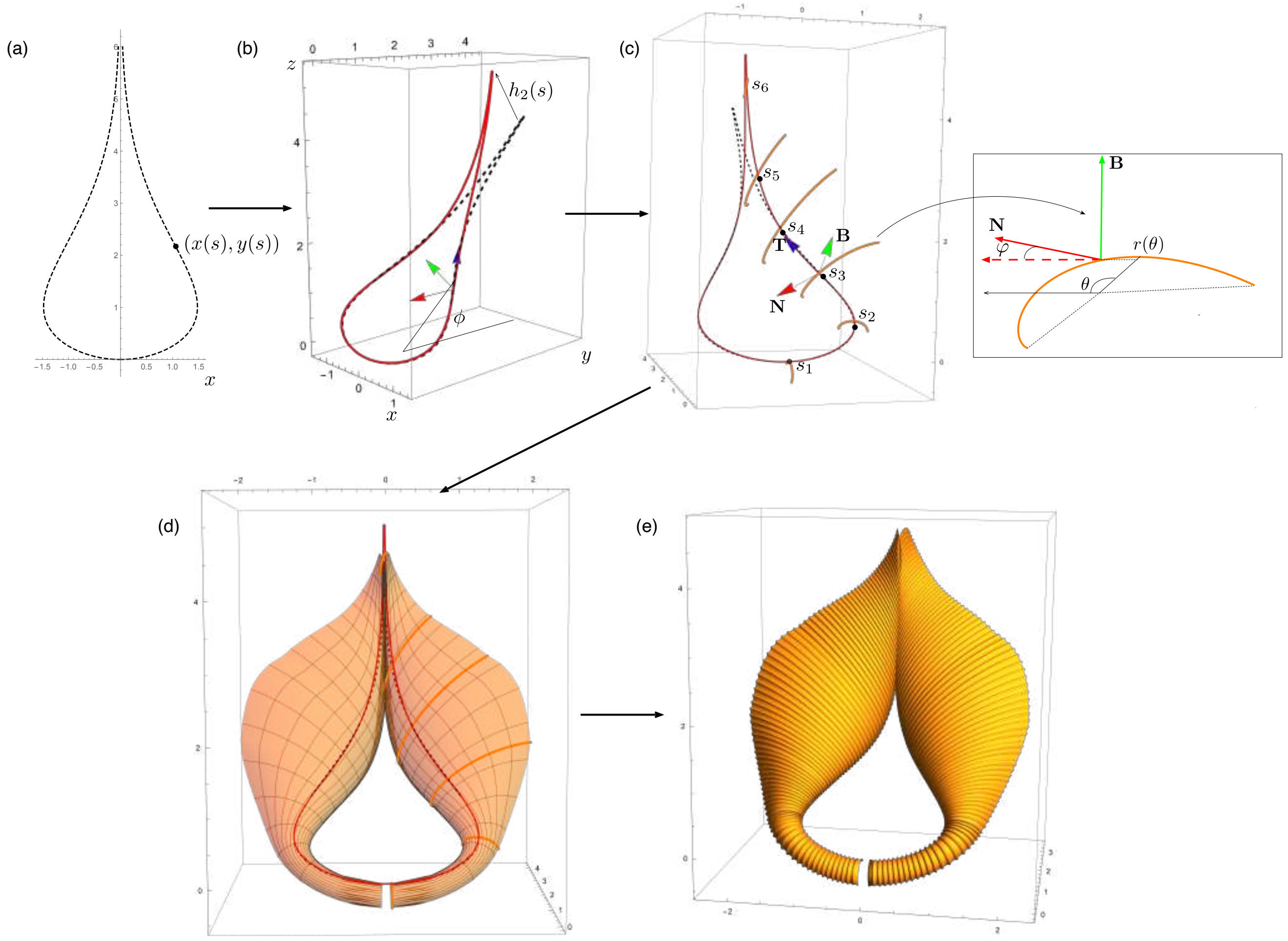

**Fig. S1.** Peristome surface construction. (a) A plane curve $(x(s), y(s))$ is constructed, and (b) translated out of plane by an amount $h_2(s)$ and rotated by angle $\phi$ about the $x$-axis, with the tangent-normal-binormal frame (TNB) is defined at each point on the curve. (c) At a discrete set of points $\{s_1, s_2, \ldots, s_n\}$, a cross-sectional curve is defined, originally in the normal-binormal plane, but rotated by angle $\varphi$ about the binormal. (d) a surface is created by interpolating the functions defining the discrete cross-sectional shapes. (e) additional features such as ribbing are added.

**B. Dry friction.** The criterion for stability under dry friction is

$$\frac{F}{N} < \mu, \qquad [8]$$

where $F$ and $N$ are the components of the reaction force $\mathbf{r}$ tangential to and normal to the surface, respectively, and $\mu$ is the friction coefficient (2). As the only force acting on the point mass is gravity, which is defined in the $z$-direction (and can be defined with unit magnitude without loss of generality), the reaction force satisfies $\mathbf{r} = \mathbf{e}_z$. The component of $\mathbf{r}$ normal to the surface has magnitude $N = |\mathbf{e}_z \cdot \mathbf{n}|$, where $\mathbf{n}$ is a unit normal vector, which can be computed via

$$\mathbf{n} = \frac{\frac{\partial \mathbf{P}}{\partial s} \times \frac{\partial \mathbf{P}}{\partial \theta}}{\left\| \frac{\partial \mathbf{P}}{\partial s} \times \frac{\partial \mathbf{P}}{\partial \theta} \right\|}. \qquad [9]$$

The frictional component of $\mathbf{r}$ is then $\mathbf{e}_z - n\mathbf{N}$, from which we determine that the point is stable if [Eq. (8)]

$$\frac{F}{N} = \frac{\sqrt{1 - (\mathbf{e}_z \cdot \mathbf{n})^2}}{|\mathbf{e}_z \cdot \mathbf{n}|} < \mu. \qquad [10]$$

**Computational aside.** It is worthwhile to note that the expressions for $G_{ij}$ as well as $\mathbf{n}$, while straightforward to compute, are very long for the surface defined by Eq. (6), and can be cumbersome to apply. For computational ease, it is useful to take advantage of the orthonormality of the right-handed basis $\{\mathbf{T}, \mathbf{N}, \mathbf{B}\}$, which satisfies the Frenet equations

$$\begin{aligned} \mathbf{T}'(s) &= \alpha(s)\kappa(s)\mathbf{N} \\ \mathbf{N}'(s) &= -\alpha(s)\kappa(s)\mathbf{T} \\ \mathbf{B}'(s) &= 0, \end{aligned} \qquad [11]$$

where the curvature $\kappa$ is given by

$$\kappa(s) = \frac{x'(s)y''(s) - x''(s)y'(s)}{\alpha(s)^3}.$$

Since the surface $\mathbf{P}$ is expressed in this basis, the scalar and vector products of the partial derivatives with respect to $s$ and $\theta$ take a reduced form when all derivatives are expressed in the basis itself using Eq. (11). Our approach was to pre-compute generic analytical expressions for the $G_{ij}$ and $\mathbf{n}$ from Eq. (6) before explicitly defining the interpolating functions. Specific computed formulas are available in the Mathematica notebooks accompanying this manuscript.

**C. Dynamic motion.** The equations of motion for a point mass sliding on the surface are determined via the Euler-Lagrange equations with energy function having components for both kinetic energy and gravitational potential energy[‡]. Noting that the path along the surface is defined by a curve in parameter space $(s(t), \theta(t))$, the position vector for the material point is given by $\mathbf{p}(t) = \mathbf{P}(s(t), \theta(t))$. The kinetic energy for mass $m$ then takes the usual form

$$\mathcal{T} = \frac{1}{2}m|\dot{\mathbf{p}}|^2, \qquad [12]$$

where overdot denotes a material derivative with respect to time, i.e. $\dot{\mathbf{p}}(t) = \dot{s}(t)\,\partial\mathbf{P}/\partial s(s(t), \theta(t)) + \dot{\theta}(t)\,\partial\mathbf{P}/\partial\theta(s(t), \theta(t))$, while the potential energy is given by

$$\mathcal{V} = mg\mathbf{p} \cdot \mathbf{e}_z, \qquad [13]$$

with $g$ the gravity of Earth. Each of these may be expressed in terms of the functions $s(t)$ and $\theta(t)$ via the parameterization given by Eq. (6), from which we may write the Lagrangian $\mathcal{L} = \mathcal{T} - \mathcal{V}$:

$$\mathcal{L}(s(t), \theta(t), \dot{s}(t), \dot{\theta}(t)) = \frac{1}{2}m|\dot{\mathbf{p}}(t)|^2 - mg\mathbf{p}(t) \cdot \mathbf{e}_z. \qquad [14]$$

The equations of motion are then given by

$$\frac{\mathrm{d}}{\mathrm{dt}}\frac{\partial\mathcal{L}}{\partial\dot{s}} - \frac{\partial\mathcal{L}}{\partial s} = 0, \qquad [15]$$

$$\frac{\mathrm{d}}{\mathrm{dt}}\frac{\partial\mathcal{L}}{\partial\dot{\theta}} - \frac{\partial\mathcal{L}}{\partial\theta} = 0. \qquad [16]$$

As initial conditions, we prescribe zero initial velocity, and choose $(s(0), \theta(0))$ to correspond to the values of seed points as described above. Eqs. (15) and (16) define two highly nonlinear coupled second order differential equations for $(s(t), \theta(t))$. Again, the calculation is straightforward; the challenge is in dealing efficiently with the cumbersome formulas. As above, this is rendered easier by expressing all derivatives back in terms of the orthonormal basis $\{\mathbf{T}, \mathbf{N}, \mathbf{B}\}$. Having reduced the formulas as outlined, the system of equations can generally be integrated forward in time in less than a second for points with simple motions (e.g. a point near the edge), or maximally within tens of seconds for more complicated motions (e.g. a point situated next to a large tooth). These were integrated using the numerical solver *NDSolve* in Mathematica,

[‡] For simplicity we neglect kinetic friction, which should be largely irrelevant with regards to the question of whether the point mass slides into or out of the pitcher.

with a stopping criterion based on reaching the edge of the surface. In particular, if $\theta(t)$ reaches $\theta_1$, the point has reached the inside edge, and we deem the mass to have fallen in the pitcher; conversely, if $\theta(t)$ reaches $\theta_2$, the point has reached the outside edge, and we deem the mass to have fallen out of the pitcher. Given the mirror symmetry, it is also possible for the point to reach the proximal point, defined by $s(t)$ reaching 0; in this event we simply reflect the velocities as appropriate and continue the integration.

The gravitational constant $g$ only has the effect of increasing or decreasing the rate with which the mass slides off the surface, and could be set arbitrarily. Here, we note that when computing the dynamic motion, we assume that the surface is fully wetted. In this case, every point is unstable except for points at which the surface is flat with respect to gravity and with appropriate sign of the mean curvature, i.e. a point situated at a local minimum. For the geometries we considered, this was not an issue, i.e. there were no attracting regions on the surface into which a point could slide into and potentially remain. If such regions did exist, the value of the gravitational constant would be important in determining the velocity with which a point entered such a region and therefore whether it became stuck; but this was not the case in our simulations.

**Fall in/fall out percentages and surface area.** Having computed the dynamic trajectories of each of the seed points, we can approximate the percentage of the surface area for which dynamic motion leads to falling in versus out by simply counting the number of seed points that lead to the different stopping criteria. That is, if the dynamic trajectories of $N_1$ seed points ended with $\theta = \theta_1$ (falling in), and there were $M$ total seed points, we approximate $N_1/M \times 100$ as the percentage of the surface for which prey "fall-in".

In terms of actual surface area, which is relevant in our energy analysis, we may compute the total surface area in the usual way, in terms of the metric tensor:

$$\mathcal{A} = \int_{s_1}^{s_n} \int_{\theta_1(s)}^{\theta_2(s)} \sqrt{\det \mathbf{G}}\, \mathrm{d}\theta \mathrm{d}s. \quad [17]$$

#### 3. Finite-sized prey

We detail the rigid body model used to investigate the effect of prey size.

**A. General setup.** We model the prey as a bilaterally symmetric rigid body with center of mass $G$ and (arbitrary) mass $m$ [Fig. S2(a)]. The body is in contact with the peristome at two points located at the same distance $\rho$ from $G$, respectively noted $C_1$ and $C_2$ (with $C_1$ the highest point). We introduce $\gamma$, the radius of gyration of the rigid body, and the angle $\alpha := 1/2 \times \widehat{C_1GC_2}$. The peristome is modeled as a circle with radius $R \equiv 1$, used as a reference length unit. The prey position is parameterized by the angle $\theta$, the angle between the prey and the vertical. We assume that the prey is subject to its own weight only. Lastly, we also prevent the leg axes from intersecting the surface by enforcing the non-penetration constraint

$$\rho \sin\alpha \le \arctan\left(\frac{\pi}{2} - \alpha\right). \quad [18]$$

**B. Mechanics.** To characterize the frictional stability of the prey, we use Erdmann's theory of Coulombian friction with multiple contact points which extends the classic notion of *friction cone* to some appropriately-defined configuration space (3, 4). Indeed, in contrast to a material point, a rigid body can be subject to both a resultant and a torque applied at $G$, that overall form a *generalized force* $(F_x, F_y, \tau) \in \mathbb{R}^3$. In particular, the *generalized friction cone* is a three-dimensional, polyhedral cone that contains all the possible generalized forces that the surface can provide under static friction while maintaining contact at both contact points. The theoretical basis of the configuration space friction cone is mathematically sophisticated; we refer the reader to Refs. (3, 4) for details. Here, we adapt the theory to our problem and briefly outline the construction of our slipping criterion. The strategy is as follows: i) construct the individual friction cone associated with each contact point; ii) obtain the composite, generalized friction cone (accounting for both contact points) by the superposition principle.

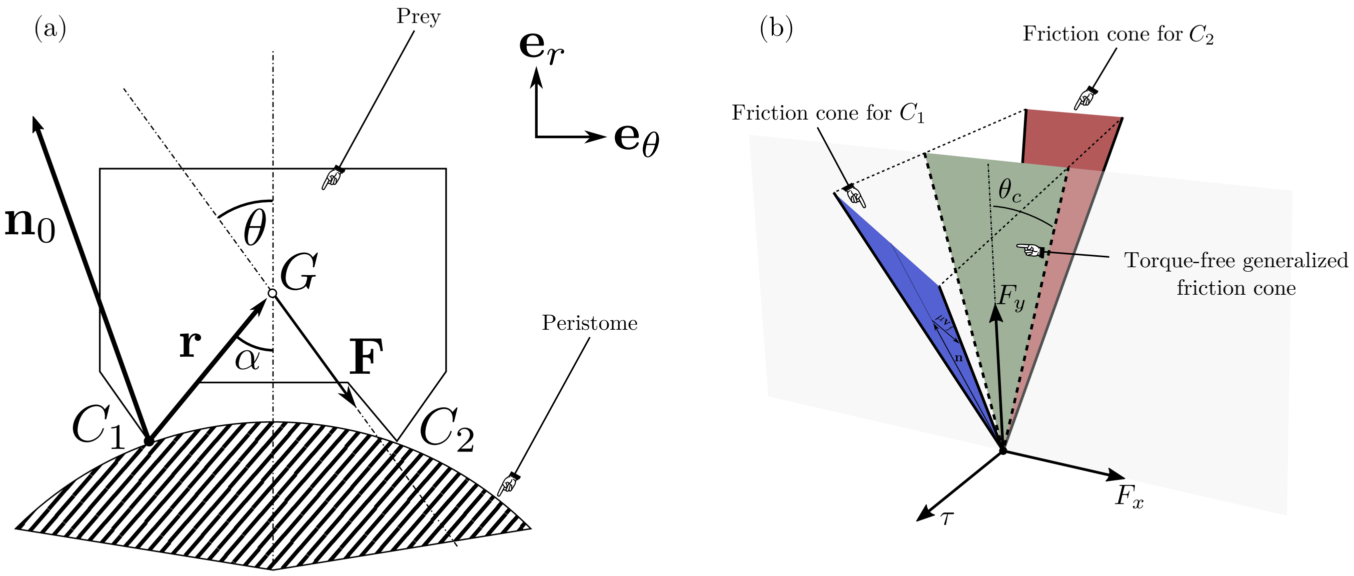

**Fig. S2.** (a) Schematic of a prey modeled as a symmetric rigid body sitting on a circular peristome. (b) The configuration space friction cone intersected with the plane $\tau = 0$ provides the critical inclination angle $\theta_c$.

We here build the friction cone for the first contact point $C_1$ (the second friction cone for $C_2$ is obtained similarly). We first define the position vector $\mathbf{r} := \overrightarrow{C_1G} = (r_x, r_y, 0) = \rho(\sin\alpha, \cos\alpha, 0)$, expressed in the canonical basis $(\mathbf{e}_\theta, \mathbf{e}_r, \mathbf{e}_\theta \times \mathbf{e}_r)$ attached to the rigid body [Fig. S2(a)]. In this basis, the weight is expressed as $\mathbf{F} = mg(\sin\theta, -\cos\theta, 0)$. Elementary geometry also provides the normal to the surface at the contact point:

$$\mathbf{n}_0 = (n_x, n_y, n_z) = \left(-\rho\sin\alpha, \sqrt{1-\rho^2\sin^2\alpha}, 0\right). \qquad [19]$$

The generalized normal follows as:

$$\mathbf{n} = \frac{1}{\Delta_n}\left(n_x, n_y, \frac{n_x r_y - n_y r_x}{\gamma}\right), \qquad [20]$$

with $\Delta_n = \sqrt{1 + (n_x r_y - n_y r_x)^2/\gamma^2}$, such that $\|\mathbf{n}\| = 1$. We stress that, while $\mathbf{n}_0$ and $\mathbf{n}$ have the same dimension, the former is a vector in the physical space, while the latter lives in the 3D configuration space. The last component of $\mathbf{n}$ corresponds to the torque about the reference point $G$, due to a unit reaction force applied at the contact point. Friction acts along the tangent through the point of contact, associated similarly with the configuration vector

$$\mathbf{v}_f = \left(n_y, -n_x, \frac{n_x r_x + n_y r_y}{\gamma}\right). \qquad [21]$$

The friction cone for $C_1$ [blue cone in Fig. S2(b)] can be written as $\{a(\Delta_n\mathbf{n} + s\mu\mathbf{v}_f) \,|\, a \geq 0, s \in [-1, +1]\}$ (the one for $C_2$ is obtained analogously). Physically, the fact that the friction cone has a component in the $\tau$ direction accounts for the fact that, if only one contact point is considered, equilibrium of the rigid body requires a non-zero applied torque to balance the applied resultant that tends to induce rotation about the contact point. Finally, the generalized friction cone, which accounts for both contact points, is obtained by the superposition principle and is given mathematically by the vector sum of the two friction cones associated with all contact points [Fig. S2(b)]. Note that to compute the friction cone, one need not compute explicitly the reaction forces at the contact points.

In our scenario, we consider a prey subject to its own weight, with no applied torque. Therefore, we consider the intersection of the three-dimensional friction cone previously constructed, with the plane $\tau = 0$ that describes the space of resultant forces, i.e. torque-free generalized forces. This intersection is a planar

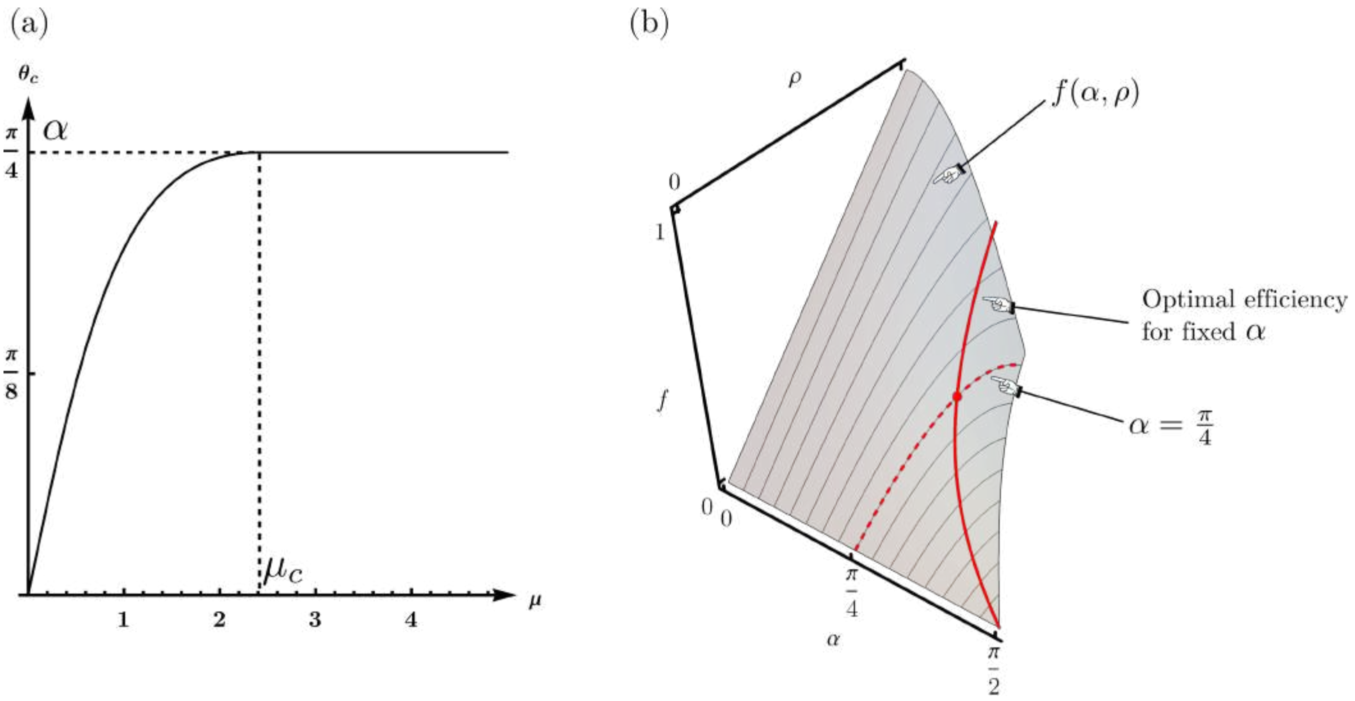

**Fig. S3.** (a) Plot of $\theta_c$ vs $\mu$, for $\alpha = \pi/4$ and for an optimal prey size, i.e. with $\rho = \rho^*(\pi/4)$ [Eq. (25)]. For $\mu > \mu_c$ [Eq. (23)], capture occurs via tumbling. (b) The efficiency function $f(\alpha, \rho)$. Maximal gain corresponds to minimal stability zone $\theta_c$, relative to the point-mass case $\theta_c \approx \mu \ll 1$.

cone [in green in Fig. S2(b)] which, as in the case of a material point, provides the maximum inclination above which slippage occurs. The director lines for this cone [thick dashed lines in Fig. S2(b)] are given by $\{a(s \sin\theta_c, \cos\theta_c, 0) \,|\, a \geq 0, s = \pm 1\}$, where $\theta_c$ denotes the critical slippage angle. Slightly tedious, but straightforward calculation of the intersection provides

$$\theta_c = \arctan\left(\frac{\mu}{1 + \rho\,(\mu^2 + 1)\left(\cos\alpha\sqrt{1 - \rho^2 \sin^2\alpha} - \sin^2\alpha\right)}\right), \qquad [22]$$

for a small enough friction coefficient $\mu \leq \mu_c$, with

$$\mu_c^2 = \frac{\sin\alpha + \rho \sin 2\alpha\sqrt{1 - \rho^2\sin^2\alpha} + \rho^2 \sin\alpha\cos 2\alpha}{\cos\alpha\,\mathrm{cotan}\,\alpha - \rho\sin 2\alpha\sqrt{1 - \rho^2\sin^2\alpha} - \rho^2\sin\alpha\cos 2\alpha}; \qquad [23]$$

otherwise

$$\theta_c = \alpha \qquad [24]$$

[Fig. S3(a)]. We first remark from Eq. (22) that $\theta_c$ is independent of the radius of gyration $\gamma$. Secondly, as expected, in the limit $\rho \to 0$ (with $\theta \leq \alpha$), we also retrieve the classic result of point-mass Coulombian friction, namely $\tan\theta_c = \mu$ (2). The latter happens to correspond to the worst-case scenario (from the plant's point of view), i.e. maximal $\theta_c$. The second case [Eq. (24)] corresponds to the situation where the reaction force at one contact point becomes negative, i.e. contact is lost and the prey tumbles into the traps (in reality, arthropod pads have some degree of adhesion, but we ignore this aspect for simplicity, arguing that adhesion is largely suppressed by the wetting of the peristome). For large friction coefficients $\mu > \mu_c$ (given $\rho$ and $\alpha$), slipping becomes impossible and capture may only occur via tumbling. In summary, a prey $(\alpha, \rho)$ satisfying the non-penetration constraint Eq. (18) will be in equilibrium if $\theta \leq \theta_c$, with $\theta_c$ described by Eqs. (22) to (24).

**C. Optimal prey size.** To explore the functional role of peristome size, we fix the size of the peristome and we look for the typical size of the most unstable prey; in other words, we consider the value of $\rho$ that minimizes $\theta_c$ (for any fixed angle $\alpha$). Remarkably, we show that this minimum is attained at a *finite* value of $\rho$, given by

$$\rho^*(\alpha) = \sqrt{\frac{\csc\alpha(\csc\alpha - 1)}{2}}, \tag{25}$$

which, interestingly, is independent of $\mu$ (only the value of the minimum will depend on $\mu$). For a slippery peristome ($\mu \ll 1$), the leading order expansion of Eq. (22), namely

$$\theta_c = \frac{\mu}{1 + \rho\left(\cos\alpha\sqrt{1 - \rho^2\sin^2\alpha} - \sin^2\alpha\right)} + \mathcal{O}\left(\mu^3\right), \tag{26}$$

further shows that, to second order in $\mu$, the relative gain in capture efficiency $\theta_c/\mu$, with respect to the worst case $\rho \to 0$, is only dependent on the geometry, via the fundamental efficiency function

$$f(\rho, \alpha) = \rho\left(\cos\alpha\sqrt{1 - \rho^2\sin^2\alpha} - \sin^2\alpha\right) \tag{27}$$

[Fig. S3(b)]. Maximum $f$ corresponds to maximum efficiency gain. The red solid line in Fig. S3(b) shows the path of maximum efficiency (for all fixed values of $\alpha$) along the surface $f(\alpha, \rho)$. For example, for a realistic angle $\alpha = \pi/4$ [red dashed line in Fig. S3(b)], the efficiency is maximal for $\rho^*(\pi/4) = \sqrt{1 - 1/\sqrt{2}} \approx 0.54$, which generates a relative efficiency gain $1 - \theta_c^*/\mu = 3 - 2\sqrt{2} \approx 17\%$ with respect to the point mass scenario, irrespective of the value of $\mu \ll 1$. Conversely, for any insect size $\rho$, the maximal capture efficiency is attained for $\alpha \to 0$. In this case, however, the insect will tumble and fall inside the pitcher before slipping.
